# A dual function of FGF signaling in *Xenopus* left-right axis formation

**DOI:** 10.1101/469791

**Authors:** Isabelle Schneider, Jennifer Kreis, Axel Schweickert, Martin Blum, Philipp Vick

## Abstract

Organ left-right (LR) asymmetry is a conserved vertebrate feature, which is regulated by left-sided activation of Nodal signaling. Nodal asymmetry is established by a leftward fluid-flow generated at the ciliated LR organizer (LRO). While the role of fibroblast growth factor (FGF) signaling pathways during mesoderm development are conserved, diverging results from different model organisms suggested a non-conserved function in LR asymmetry. Here, we demonstrate that FGF is required during gastrulation in a dual function at consecutive stages of *Xenopus* embryonic development. In the early gastrula, FGF is necessary for LRO precursor induction, acting in parallel to FGF-mediated mesoderm induction. During late gastrulation, the FGF/Ca^2+^-branch is required for specification of the flow sensing lateral LRO cells, a function related to FGF-mediated mesoderm morphogenesis. This second function in addition requires input from the calcium channel Polycystin-2. Thus, analogous to mesoderm development, FGF activity is required in a dual role for laterality specification, namely for generating and sensing of leftward flow. Moreover, our data show that FGF functions in LR asymmetric development are conserved across vertebrate species, from fish to mammals.

## Introduction

Left-right (LR) asymmetry of inner organs is present in all deuterostome lineages and depends on the left-sided activation of the evolutionarily highly conserved Nodal signaling cascade. In fish, amphibian and mammalian embryos, the symmetry breaking event, which activates this asymmetric gene cascade in the lateral plate mesoderm (LPM), is represented by a cilia-based extracellular fluid-flow at the left-right organizer (LRO) of the early neurula embryo (Dasgupta and Amack, 2016; Yoshiba and Hamada, 2014). The flow generating LRO epithelium consists of monociliated cells of mesodermal fate, which are transiently embedded in the endodermal endothelium of the forming archenteron at the posterior end of the notochord (Blum et al., 2007). In amphibians, it is of triangular shape and addressed as gastrocoel roof plate (GRP; Blum et al., 2009b). The *Xenopus* GRP is subdivided into a medial flow-generating part, with posteriorly localized motile cilia, and lateral cells with non-polarized and immotile cilia, which presumably sense flow. The former cells will later integrate into notochord and hypochord, while the latter will contribute to somites (Boskovski et al., 2013; Shook et al., 2004). Somitic GRP cells are characterized by co-expression of *nodal* and the Nodal inhibitor *dand5*. Flow is thought to downregulate *dand5*, de-repress Nodal and induce transfer of an asymmetric signal to the left LPM (Schweickert et al., 2010). The GRP derives from the superficial mesoderm (SM; Shook et al., 2004) of the early gastrula, which represents the outer-most layer of the Spemann organizer. The SM is marked by the expression of the transcription factor *forkhead box J1* (*foxj1*; Blum et al., 2014; Stubbs et al., 2008), which induces motile ciliogenesis.

A number of signaling pathways have been implicated in LR development, although conserved functions have only been described for a few, such as Nodal (Namigai et al., 2014). Conflicting results have been obtained for FGF signaling. In chick and rabbit embryos, the ligand Fgf8 seems to act as a right determinant, repressing asymmetric gene expression in the right LPM (Boettger et al., 1999; Feistel and Blum, 2008; Fischer et al., 2002). In the mouse, in contrast, Fgf8 acts as a left determinant. Here, FGF signaling is required both for *nodal* gene expression at the LRO and for asymmetric gene expression in the LPM (Meyers and Martin, 1999; Oki et al., 2010). In addition, FGF signaling has been implicated as regulator of vesicle release at the LRO, which transfer through flow to the left side, where they activate Nodal signaling (Tanaka et al., 2005). In teleost fish, FGF signals were shown to be required for symmetry breakage upstream of leftward flow. Loss of FGF signaling resulted in reduction or loss of LRO-cilia and, consequently, LR defects (Hong and Dawid, 2009; Neugebauer et al., 2009; Yamauchi et al., 2009). Reduced ciliary lengths were reported in the *Xenopus* GRP, but GRP morphogenesis, *foxj1*, flow and Nodal cascade genes have not been analyzed (Neugebauer et al., 2009). Thus, the role of FGF signaling seems to vary in the different model organisms analyzed to date.

Here we studied the role of FGF signaling in LR axis formation in *Xenopus*. The frog is accessible to manipulation and analysis of all stages of LR development, from SM specification to GRP morphogenesis, leftward flow, Nodal cascade induction and organ morphogenesis (Blum et al., 2014; Blum et al., 2009a). Through pharmacological and molecular manipulation of Fgfr1 signaling, we identified a dual role of FGF signaling. During early gastrula, FGF was necessary for induction of the SM upstream of *foxj1*. During late gastrulation, a second, cilia-independent function was involved in specification of the LRO flow sensor, downstream of *foxj1* induction. This late phenotype was caused by inhibition of FGF/Ca^2+^-signaling, a function which required Polycystin-2 as well. Our data reconcile conflicting findings in model organisms and demonstrate a conserved role of FGF signaling in LR axis formation, in much the same dual way as has been shown for mesoderm induction and morphogenesis.

## Results

### FGF signaling is necessary for LR asymmetry during gastrulation

In order to determine the critical time frame of FGF signaling in *Xenopus* LR development, we treated embryos of defined stages with SU5402, an inhibitor of the tyrosine kinase activity of Fgfr1 (Fig. 1A). SU5402 incubation at late blastula or early gastrula stages (until st. 10.5) impaired gastrulation movements leading to blastopore closure defects, reduced somitic expression of *myogenic differentiation 1* (*myod1*), and shortened anterior-posterior axis, as previously described for inhibition of FGF signaling (Fig. 1B; Amaya et al., 1991). Mesodermal defects could be minimized by treatment at mid gastrulation (st.11/11.5), and circumvented completely when treated from late gastrula stages onwards (st.12/12.5; Fig. 1B and data not shown). In all cases, however, i.e. from early, mid or late gastrulation onwards, FGF-inhibition resulted in absence of *pitx2c* expression in the left LPM (Fig. 1A-C). Treatments at later stages did not impact on laterality (Fig. 1C). Inhibition of FGF signaling did also not impair the competence of LPM tissue to respond to Nodal, as injection of a *nodal1*-DNA construct into the LPM of SU5402-treated embryos rescued *pitx2c* expression (Fig. 1C). These experiments revealed a sensitive time window for FGF signaling in LR axis development from late blastula until late gastrulation, i.e. before establishment of leftward flow and thus, upstream of *nodal1* activation in the left LPM.

**Fig. 1.**
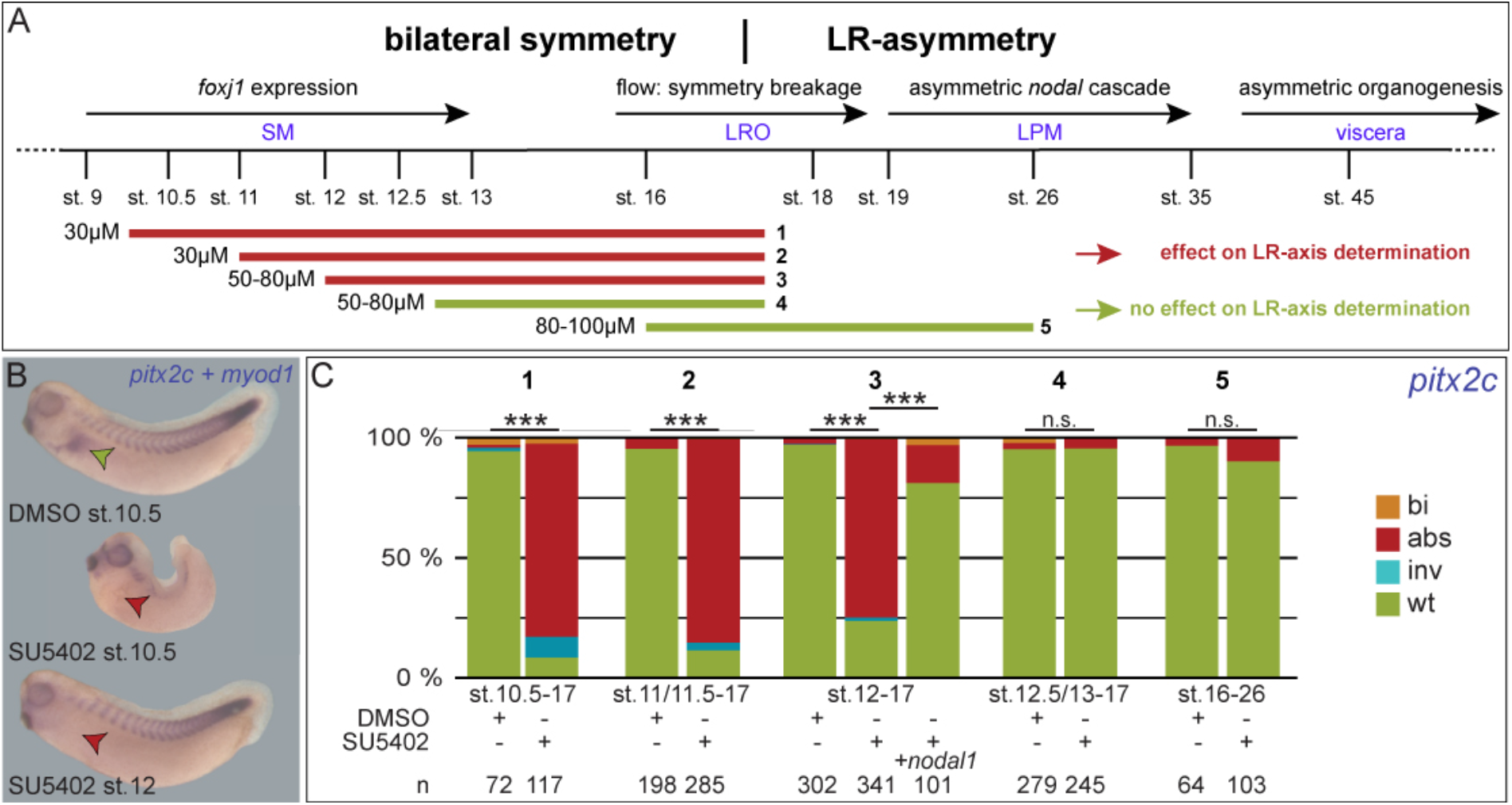
FGF signaling is required for LR axis development during gastrulation. (A) Timeline of events relevant for LR axis development (top), and SU5402 incubation periods (bottom; marked as 1-5). (B) Representative specimens encountered following co-staining for *pitx2c* and *myod1* mRNA expression. Top, DMSO-treated embryo; wildtype morphology and expression patterns (left-sided expression of *pitx2c*, marked by green arrowhead, and of *myod1* in somites. Middle: specimen treated with SU5402 at early gastrulation (stage 10.5) displaying lack of *pitx2c* expression (red arrowhead). Bottom: specimen treated with SU5402 at late gastrula (stage 12). Please note strongly decreased expression of *myod* upon early, but not late SU5402 treatments. (C) Quantification of *pitx2c* expression patterns in specimens treated during indicated time periods (1-5). abs, absent; bi, bilateral; inv, inverse; LPM, lateral plate mesoderm; LRO, left-right organizer; n.s., not significant; SM, superficial mesoderm; st., stage; wt, wildtype; n are number of embryos analyzed.

### A first FGF signal is required for *foxj1* induction during early gastrulation

Loss of asymmetric *pitx2c* expression could be caused by lack of symmetry breakage due to impaired leftward flow. Gastrulation defects, however, prevented the analysis of leftward flow in embryos treated until stage 10.5 (Fig. 1B). We therefore decided to analyze the specification of the precursor tissue from which the LRO derives, namely the superficial mesoderm (SM). This reasoning was further supported by our recent demonstration that knockdown of the Fgfr1 ligand *nodal3.1* (*nodal homolog 3, gene 1*; previously known as *xnr3*) in the SM inhibited *foxj1* induction (Vick et al., 2018; Yokota et al., 2003). In agreement with this notion, embryos treated with SU5402 at early gastrula (stage 10.5) showed a significant reduction of *foxj1* in the SM, confirming a role of Nodal3.1/FGF in setting-up a functional GRP (Fig. 2A-C). Injection of a dominant-negative version of Fgfr1 (*dnfgfr1*; Amaya et al., 1991) mimicked SU5402 treatment and resulted in a significant loss of *foxj1* expression as well (Fig. S1A, B). Together, these experiments demonstrated that an early FGF signal mediated through Fgfr1 was required for parallel induction of gastrulation movements and *foxj1* expression in the SM.

**Fig. 2.**
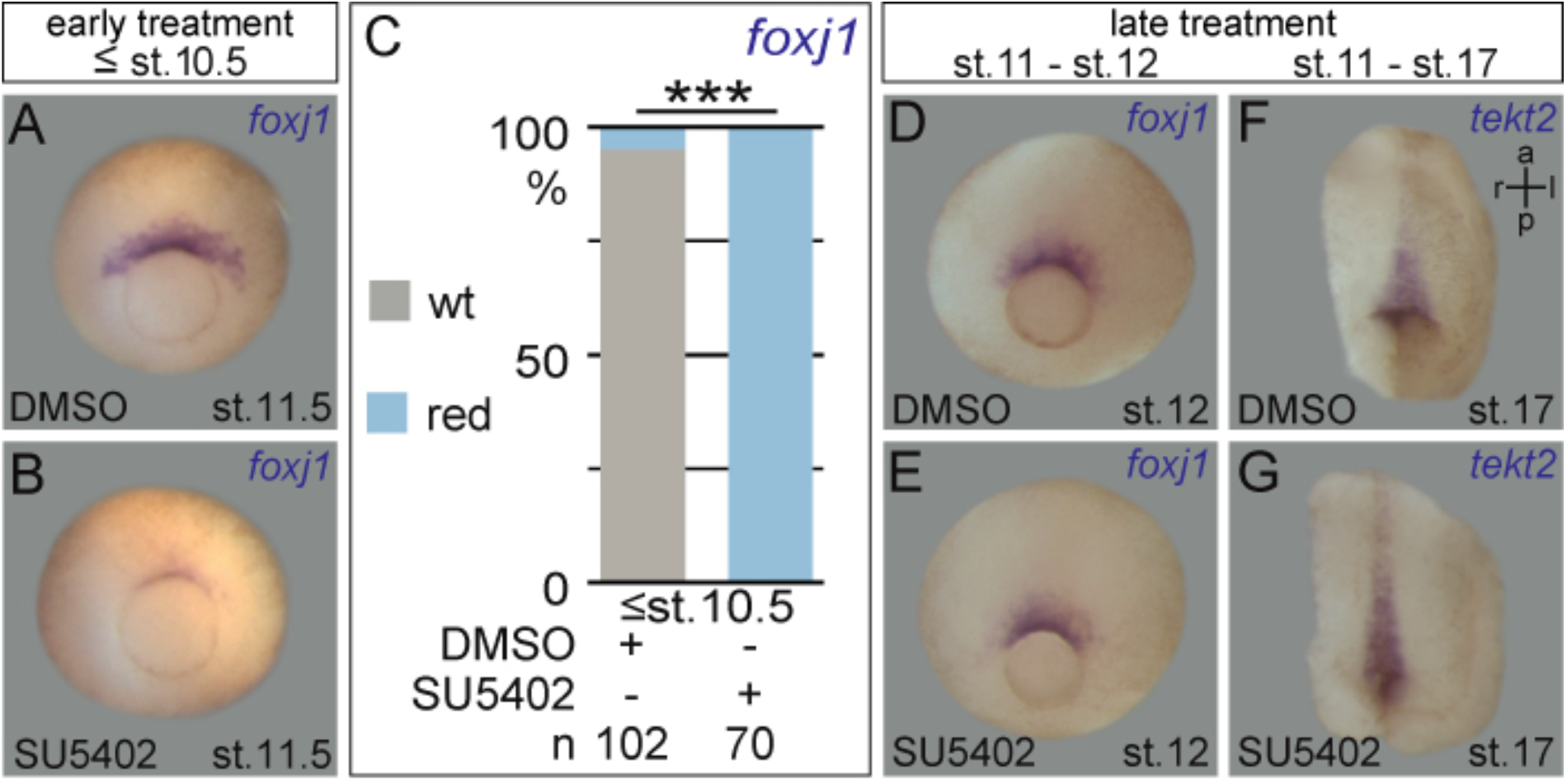
Inhibition of FGF signaling at early gastrulation acts upstream of *foxj1*. (A-C) Downregulation of SM marker gene *foxj1* in embryos treated with SU5402 up to gastrula stage 10.5. (A, B) Representative *foxj1* expression patterns at stage 11.5 in embryo treated with DMSO (A) or SU5402 (B). (C) Quantification of results. (D-G) Unaltered expression of *foxj1* (in the SM) and *tekt2* (in the GRP) of embryos treated with SU5402 from mid gastrula stages onward. Representative examples of expression patterns of *foxj1* (D, E), and *tekt2* (F, G) in embryos treated with DMSO (D, F) or 80µM SU5402 (E, G). Embryos in (F) and (D) are ventral views of dorsal explants displaying archenteron roof and embedded GRP. a, anterior; l, left; p, posterior; r, right; red, reduced; st., stage; wt, wildtype; n, number of embryos analyzed.

### A second FGF signal regulates Nodal cascade induction independent of *foxj1* and motile cilia

Interestingly, SU5402 incubation at late gastrula stages had no impact on *foxj1*, in agreement with the robust expression of *foxj1* at these stages (Fig. 2D, E; cf. Beyer et al., 2012; Stubbs et al., 2008). Injection of low doses of *dnfgfr1* did not cause a significant loss of *foxj1* expression either, nor did it block gastrulation movements (Fig. S1C and data not shown). At neurula stages, expression of the Foxj1 target gene *tekt2* in the GRP was apparently unaffected by late gastrula stage SU5402 incubations. The *tekt2* expression domain, however, appeared narrower than that of control embryos (Fig. 2F, G). We therefore analyzed cilia formation and functionality by scanning electron microscopy (SEM) and flow analysis, to test for effects downstream of ciliary gene activation. SEM analysis did not reveal significant differences in cilia length, GRP ciliation or posterior polarization of cilia between treated and control specimens (Fig. S1D-H). Low dose *dnfgfr1* injections did not alter ciliary length either (Fig. S1I). Cilia motility and leftward flow were also unaffected, as demonstrated by flow analysis of wildtype and SU6402-treated GRP explants from stage 17/18 embryos (Fig. S1J-L; (Schweickert et al., 2007). Together, these data hint at two different modes of action of FGF signaling in *Xenopus* LR axis development: a cilia- and *foxj1*-dependent function at early gastrulation, and a later, cilia-independent mode of action downstream of flow and upstream of *nodal1* expression in the LPM.

### Late FGF signaling is necessary and sufficient for LRO flow sensor formation

Next, we asked if signaling downstream of flow was impaired in embryos treated at late gastrula stages, i.e. whether the left-asymmetric flow signal was perceived and transferred to the LPM. In wild-type embryos, polarized motile cilia at the center of the GRP generate flow which is perceived by immotile and central cilia on lateral GRP cells (Boskovski et al., 2013; Schweickert et al., 2007; Shook et al., 2004). Sensory cells are characterized by the co-expression of both *nodal1* and its inhibitor *dand5* (Schweickert et al., 2010; Vonica and Brivanlou, 2007). SU5402 incubation of stage 12 resulted in absence of both *nodal1* and *dand5* expression (Fig. 3A, B, D, E, G), while effects were greatly reduced upon slightly later incubations at stage 13 (Fig. 3C, F, G). Low-dose *dnfgfr1* injections in a sided manner resulted in a significant reduction or loss of *nodal1* and *dand5* on the injected side (Fig. S2). In line with this notion, activation of FGF signaling by injection of the ligand *fgf8b* into the lateral GRP was sufficient to increase the expression of both *nodal1* and *dand5* in a highly significant proportion of specimens (Fig. 3H-L). Overexpression was achieved through injections of a DNA construct, which gets only transcribed after activation of the zygotic genome at late blastula stages, in order to prevent earlier FGF-induced developmental defects. Together, these experiments pointed to a function of FGF signaling during late gastrulation to promote the lateral, somitic fate of the GRP.

**Fig. 3.**
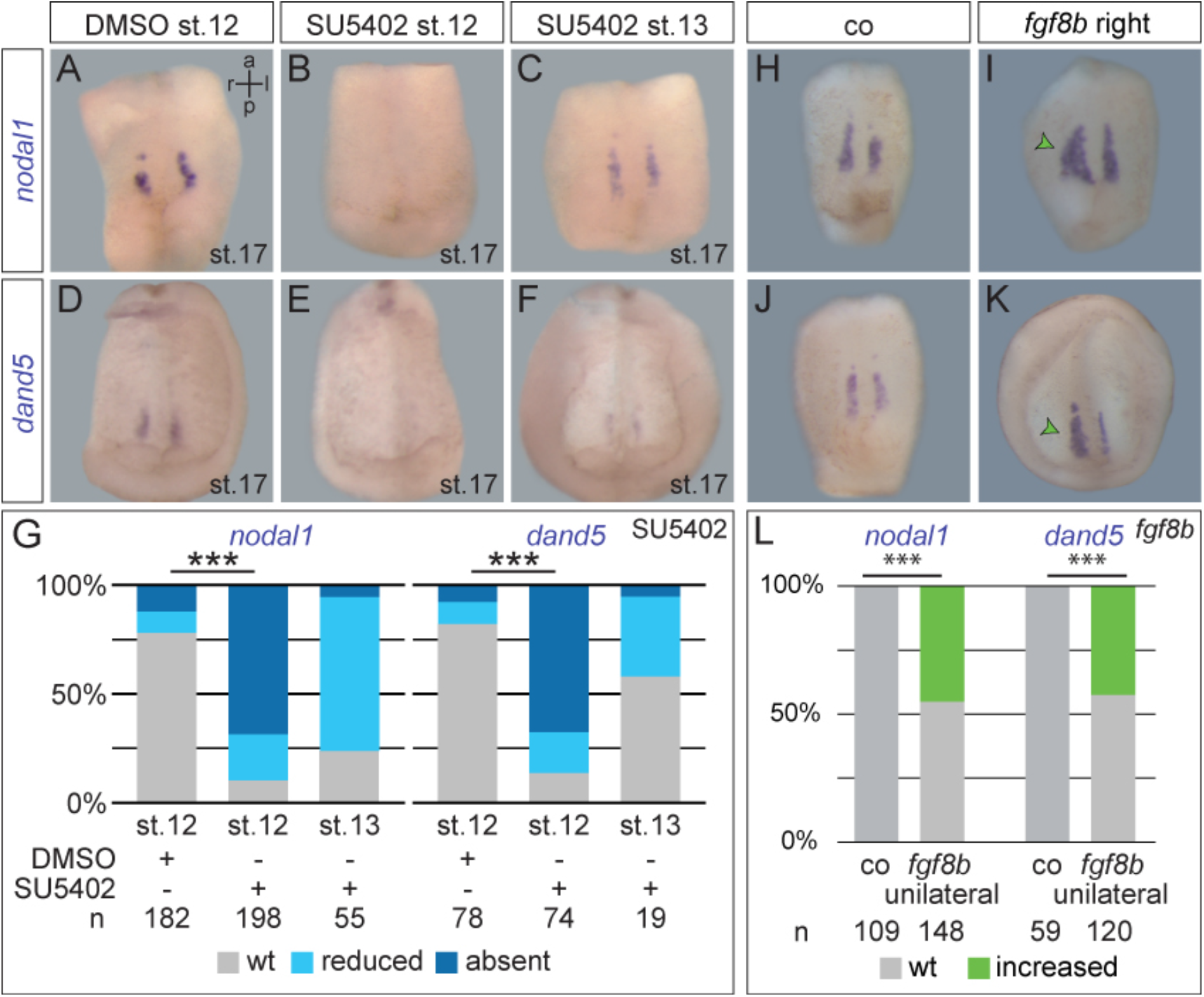
FGF signaling regulates somitic GRP specification from mid to late gastrulation. (A-F) Analysis of *nodal1* (A-C) and *dand5* (D-F) mRNA expression in st. 17 embryos treated with DMSO (A, D) or 50µM SU5402 (B, C, E, F) from st. 12 (A, B, D, E) or 13 (C, F) onwards. Lateral GRP-specific expression patterns shown in ventral view of dorsal explants. (G) Quantification of results. (H-K) Analysis of *nodal1* (H, I) and *dand5* (J, K) in st. 17 control specimens (H, J) or embryos unilaterally injected with *fgf8b* DNA (5-8pg), specifically targeting the right expression domain of *nodal1*/*dand5*. Injected embryos (I, K) showed a lateral increase of expression domains on the targeted side. (L) Quantification of results. a, anterior; l, left; n.s., not significant; p, posterior; r, right; st., stage; wt, wildtype; n are numbers of embryos analyzed.

As both *nodal1* and *dand5* expression were lost, we wondered if FGF-inhibition prevented activation of these genes, or whether the late FGF signal was required for formation of sensory cells. These cells later detach from the gastrocoel roof and integrate into the presomitic mesoderm (Shook et al., 2004). Their somitic fate is marked already in the GRP by expression of *myod1* (Fig. 4A, B, E; Schweickert et al., 2010). This contribution of *myod1*-positive cells to the GRP was essentially lost in specimens treated with SU5402-incubation at stage 12 (Fig. 4C, E), to a lesser degree upon incubation at stage 13 (Fig. 4D, E). Low doses of *dnfgfr1* injections likewise resulted in lack of *myod1*-positive lateral GRP cells on the injected side (Fig. S3A), demonstrating the specificity of effects. To assess whether this absence of *myod1* in the GRP was a failure of gene induction, or a general failure of somitic GRP cell formation, SEM analyses were performed. In control embryos, somitic GRP cells were clearly distinguishable from the central GRP, which harbor polarized cilia, and from the much larger non-ciliated lateral endodermal cells (Fig. 4F, G). Strikingly, in SU5402-treated embryos, somitic GRP cells were lost, such that the central part of the GRP was found directly adjacent to the lateral endodermal cells (Fig. 4H, I). The width of treated GRPs was narrower, encompassing just the size of a wild-type central GRP (Fig. 4J). This notion was confirmed by SEM analysis of *dnfgfr1*-injected embryos, which again showed strong reduction of the somitic part of the GRP on the injected side (Fig. S3B). Finally, unilateral injection of *fgf8b* increased the size of the somitic GRP as well (Fig. S3C). Together, these analyses demonstrated a requirement of Fgfr1-activation for sensory GRP specification during late gastrulation (Fig. 4K).

**Fig. 4.**
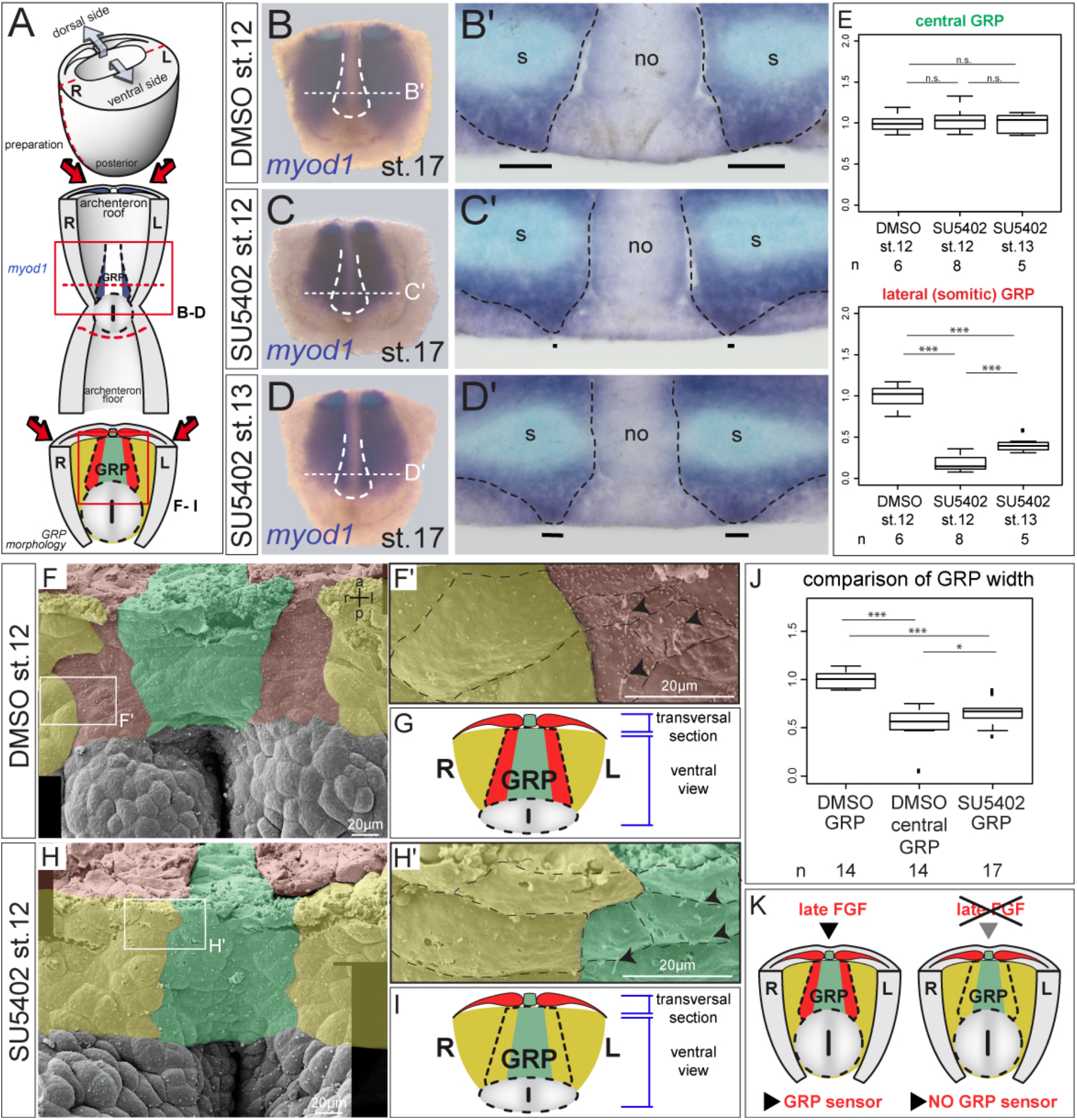
FGF-inhibition during late gastrulation impairs somitic GRP formation. (A) Schematic depiction of explant preparation and analysis. Neurula-stage embryos were bisected transversally, the posterior part bisected into dorsal and ventral halves (top). Dorsal halves were either analyzed for *myod1*-expression (B-E), or bisected transversally to visualize the GRP from top by SEM (bottom; F-J). *myod1*-expression in somites and lateral GRP indicated in blue (middle). Lateral GRP and somites indicated in red, central GRP and notochord in green, and endoderm in yellow (bottom). (B-D) ISH analysis of *myod1* expression in st. 17 embryos treated with DMSO (B) or 50µM SU5402 (C, D) from stage 12 (B, C) or stage 13 (D) onwards. Ventral view of representative dorsal explant. Position of GRP indicated by dashed line. (B’-D’) Histological analysis of *myod1* expression by transversal sections at levels indicated in (B-D). Black bars highlight width of *myod1*-positive GRP, which was strongly (C’) or slightly (D’) reduced in SU5402-treated embryos. (E) Quantification of widths of central and somitic GRP areas in DMSO- and SU5402-treated embryos. Mean width of DMSO-treated GRPs was set to 1.0. (F-I) SEM of transversally broken st. 17 dorsal explants (ventral view) treated with DMSO (F) or 60µM SU5402 (H) from st. 12 onwards. Coloring in (F, H) and in blow-ups (F’, H’) mark the central (green), and lateral somitic GRP (red), and the flanking lateral endoderm (yellow) in representative DMSO- or SU5402-treated specimens. Apical surface (F’, H’) of individual cells outlined by dashed lines. Black arrowheads highlight cilia of somitic (F’), or central GRP cells (H’). Please note lack of somitic GRP after late SU5402-treatments, as illustrated in (G) and (I). (J) Quantification of GRP widths. (K) Scheme summarizing results of late FGF-inhibition. a, anterior; GRP, gastrocoel roof plate; l, left; n.s., not significant; p, posterior; r, right; s, somite; st., stage; wt, wildtype; n are numbers of embryos analyzed.

### Sprouty-induced inhibition of FGF/Ca^2+^-signaling blocks the GRP flow sensor

A dual role of FGF signaling has been previously described in the context of mesoderm induction and subsequent morphogenesis (Nutt et al., 2001; Sivak et al., 2005). Specifically, MAPK-dependent FGF signaling is necessary during early/mid gastrulation for mesoderm induction and maintenance. At mid-gastrulation, the PKC-dependent FGF/Ca^2+^-branch is required for morphogenetic gastrulation movements, which, however, does not impact on mesoderm induction. The FGF-antagonist *sprouty* is expressed during early gastrulation; overexpression does not impact on MAPK-mediated mesoderm induction, but blocks FGF/Ca^2+^-activation during late gastrulation, i.e. specifically regulates FGF-mediated morphogenesis (Nutt et al., 2001; Sivak et al., 2005). The dual role of FGF signaling in LR axis formation made us wonder whether the FGF/Ca^2+^ branch was likewise involved in the FGF-mediated specification of the sensory GRP.

In order to test this option, we injected *spry1* mRNA to target the SM and lateral GRP cells, respectively. Sprouty overexpression did not affect *foxj1* expression in the SM, nor subsequent expression of its target *tekt2* in the GRP (Fig. 5A-F). Mesoderm induction and mesodermal marker gene activation (*brachyury*, *myod1*) were unaffected, as described (Sivak et al., 2005); data not shown). However, the width of *tekt2* expression was reduced following *spry1* injections (compare Fig. 5D and E). Expression of *nodal1* and *dand5* was lost on the injected side of unilaterally injected specimen (Fig. 5G-L). The loss of the lateral GRP cells upon *spry1* injection was confirmed by SEM analysis (data not shown). Together, these results demonstrate a requirement of the FGF/Ca^2+^-pathway for morphogenesis of the LRO flow sensor, i.e. somitic GRP cell formation during late gastrulation.

**Fig. 5.**
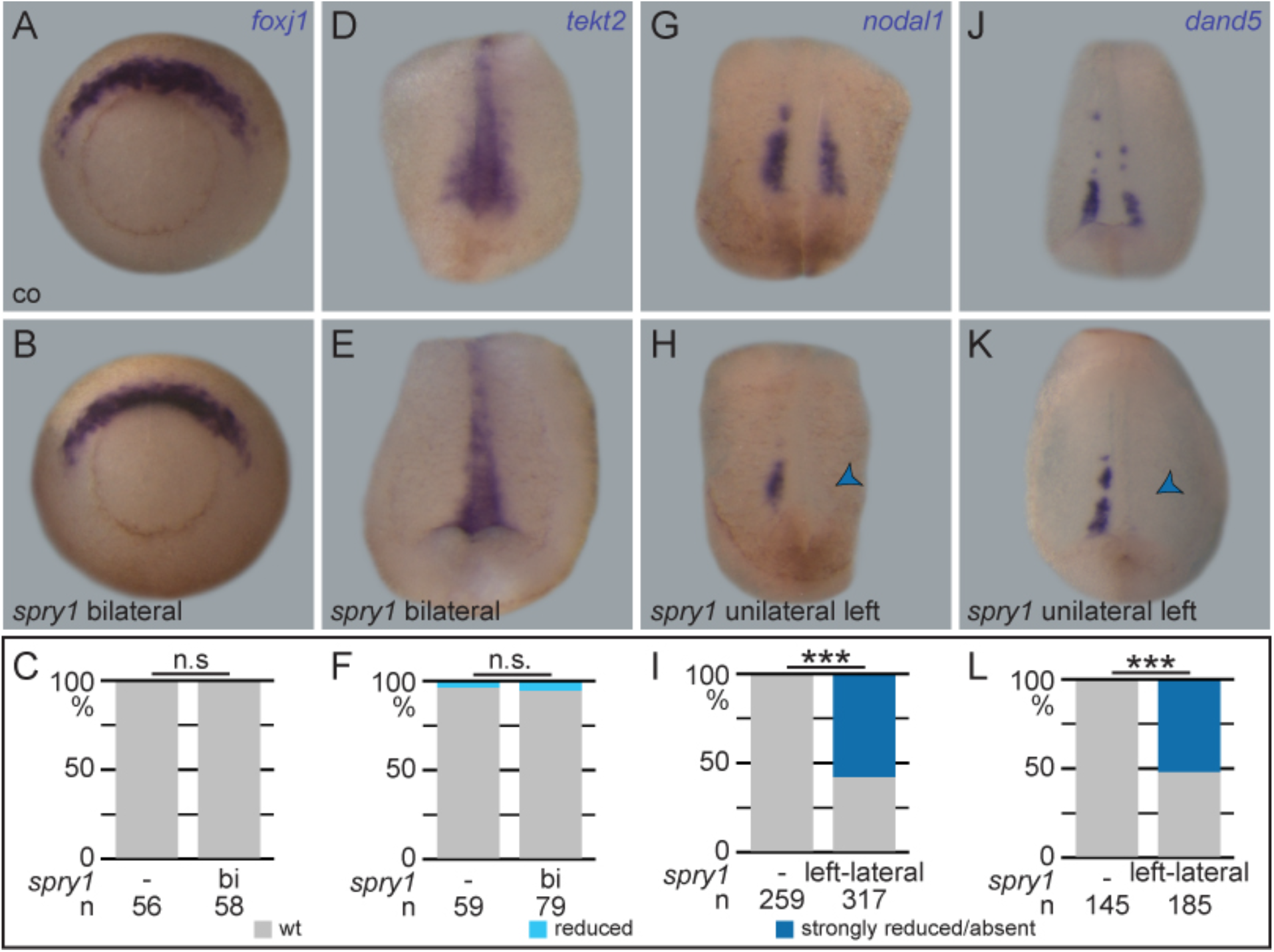
Sprouty-mediated FGF-inhibition blocks specification of somitic GRP. (A-F) ISH analysis of *foxj1* in st. 11 embryos (A-C) and *tekt2* in st. 17 embryos (D-F). (C, F, I, L) Quantification of results. (A-C) Wildtype *foxj1* expression in control specimens (A) and embryos bilaterally injected with *spry1* mRNA. (D-F) *tekt2* expression was narrowed in injected specimen (E) as compared to control (D). (G-L) *nodal1* (G-I) and *dand5* (J-L) were absent on the injected side of st. 17 embryos after unilateral injection with *spry1* mRNA (blue arrowheads). *sprouty1* mRNAs were injected at 320-800pg. (A, B) vegetal views, dorsal side up; (D, E, G, H, J, K) ventral views of dorsal explants. bi, bilateral; n.s., not significant; wt, wildtype; n, numbers of embryos analyzed.

### FGF/Ca^2+^ and *pkd2* synergize in LRO sensor formation

Like FGF, *pkd2* is required in a dual way during symmetry breakage in *Xenopus* (Vick et al., 2018). In a first stage, it cooperates with the FGF ligand *nodal3.1* to induce *foxj1* in the SM. Later, it is required for somitic GRP cells and initiation of *nodal1*/*dand5* (Vick et al., 2018). As *pkd2* encodes the calcium channel Polycystin-2 (Busch et al., 2017; Koulen et al., 2002), we reasoned that it could interact synergistically with the FGF/Ca^2+^-pathway. To analyze this possibility, we injected a characterized *pkd2* antisense morpholino oligomer (MO; Tran et al., 2010; Vick et al., 2018) to target the *nodal1*/*dand5* expressing somitic GRP cells. The MO prevented expression of both genes specifically on the injected side, as described (Fig. S4AD; cf. (Vick et al., 2018). To test for genetic interaction, we lowered both *pkd2*-MO dose and concentration of *spry1* mRNA to only mildly affect *nodal1* and *dand5* expression on the injected side (Fig. 6A-F). Co-injections of sub-phenotypic doses, however, resulted in a highly significant reduction of both genes (Fig. 6D-F). These experiments indicated that *pkd2* loss- and *sprouty* gain-of-function impact on the same process, namely somitic GRP-induction. As Sprouty presumably blocks the FGF/Ca^2+^-pathway upstream of ER-mediated intracellular Ca^2+^-release (Akbulut et al., 2010; Nutt et al., 2001), *pkd2* might be able to rescue the *spry1-*induced loss of sensory GRP cells. Indeed, co-injection of *spry1* and *pkd2* partially restored *nodal1* expression in lateral GRP cells (Fig. 6G-J). These final experiments demonstrated a synergy of *pkd2* and the FGF/Ca^2+^-branch of FGF signaling in inducing the somitic GRP, i.e. for specification of the flow sensor of the left-right organizer.

**Fig. 6.**
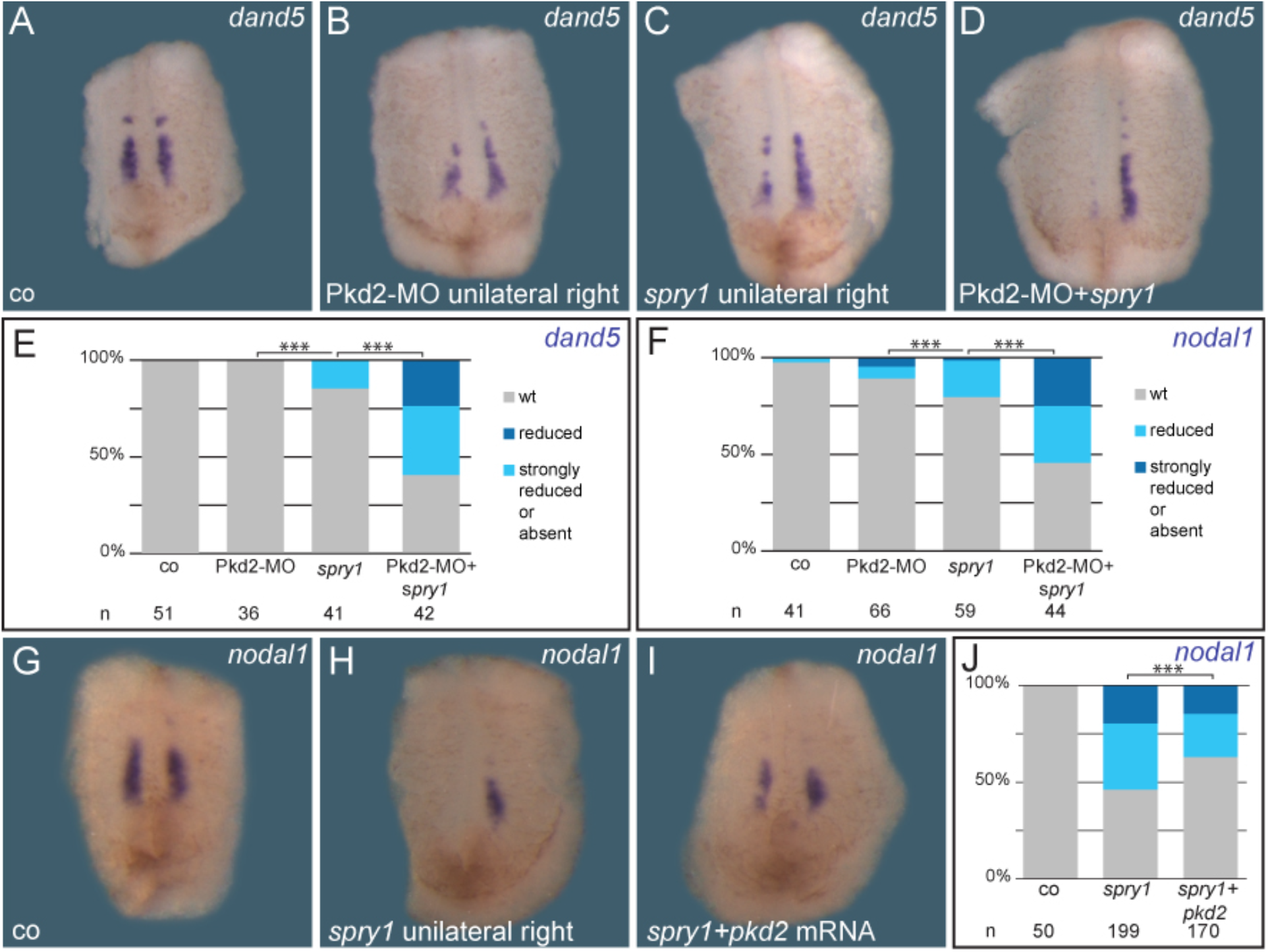
*sprouty1* and *pkd2* cooperate in somitic GRP specification. (A-F) Epistatic analysis of FGF/Ca^2+^ and Polycystin-2 function. Simultaneous overexpression of *spry1* and knockdown of *pkd2* using sub-phenotypic doses. ISH analysis of *dand5* (A-E) or *nodal1* (F) in st. 17 embryos showed wildtype expression in controls (A, E, F). No or very minor reductions were observed following unilateral injection of low doses of *spry1* mRNA (160pg; C, E, F) or Pkd2-MO (0.5pmol; B, E, F), while combinations of both yielded a high fraction (>50%) of embryos with reduced or absent gene expression levels (D-F). (G-J) Rescue of *nodal1* expression (G) in *spry1*-injected specimens (H, 320-800pg) by co-injection of *pkd2* mRNA (I). (J) Quantification of results. (G-I). Ventral views of dorsal explants. wt, wildtype; n, numbers of embryos analyzed.

## Discussion

Our analysis of FGF signaling during left-right axis formation in *Xenopus* demonstrated a dual role, namely for specification of the LRO precursor, the SM, during early gastrulation, and a later, independent role for sensory cell formation in the LRO itself. These findings reconcile conflicting data reported previously from zebrafish and mouse. Ciliogenesis of the LRO and f*oxj1* expression, which marks the LRO and its precursor in vertebrates employing cilia-mediated symmetry breaking, were either reduced or lost in zebrafish following SU5402-treatment or knockdown of *fgfr1* or *fgf8* (Caron et al., 2012; Hong and Dawid, 2009; Neugebauer et al., 2009). In contrast, data reported from a hypomorphic *fgf8* mouse line suggested that lateral, sensory LRO cells might be missing, based on the absence of *nodal* expression at the LRO, which is an evolutionarily conserved hallmark of ciliated LROs (Blum et al., 2007; Blum et al., 2009b; Meyers and Martin, 1999). Analysis of FGF signaling during late gastrulation using SU5402 also resulted in loss of *nodal* expression in the mouse LRO (Oki et al., 2010). Whether or not this function was mediated through the FGF/Ca^2+^ pathway, as shown here, has not been addressed in either of these studies, nor a possible role in sensory cell formation, a concept that emerged only later (McGrath et al., 2003; Tabin and Vogan, 2003). Finally, it may be worthwhile re-evaluating LRO specification (*foxj1* expression) and ciliation in mouse embryos with reduced FGF signaling (in the hypomorphic line or after early SU5402-treatment), in order to analyze whether the first phase of FGF-dependent laterality formation, i.e. LRO precursor formation, is conserved in mammalian embryos as well.

The combined data on FGF function in cilia-dependent LR axis formation in the vertebrates strongly suggest an evolutionarily conserved dual role during early and late gastrulation. Interestingly, this dual function parallels the known role of FGF signaling during *Xenopus* mesoderm induction (phase 1 during early gastrulation; Amaya et al., 1993; Schulte-Merker and Smith, 1995) and morphogenesis (phase 2 during late gastrulation; Dorey and Amaya, 2010; Pownall and Isaacs, 2010). Both processes are intimately linked through the timing of events. Morphogenetic gastrulation behavior in phase 2 requires calcium, and is mediated through the FGF/Ca^2+^ pathway as well (Sivak et al., 2005; Wallingford et al., 2001; Wang and Steinbeisser, 2009). In the lateral GRP of *Xenopus*, Polycystin-2 might cooperate with FGF/Ca^2+^ downstream of inositol trisphosphate receptor activation at the level of intracellular calcium release, a positive interaction that has been demonstrated in *Xenopus* oocytes (Berridge et al., 2000; Dorey and Amaya, 2010; Li et al., 2005). This notion is supported by Thapsigargin-mediated interference with calcium fluxes, which also resulted in lack of *nodal* expression in mouse and *Xenopus* LRO (Takao et al., 2013; Thastrup et al., 1990; Vick et al., 2018).

Although not considered mesodermal at the stage of emergence and function, cells at the *Xenopus* SM (i.e. superficial cells of the organizer) and LRO (embedded in the archenteron) are fated to become mesodermal: they integrate into the notochord (flow-generating central LRO cells) and somites (flow-sensing lateral LRO cells) once the LRO submerges under the endodermal cells, which close the gap in the posterior dorsal archenteron during neurulation (Shook et al., 2004). Thus, SM induction may be considered as one aspect of mesoderm induction, and LRO formation as a morphogenetic process during early mesoderm differentiation. In that sense, LRO cells take a unique detour to assume their final mesodermal fate, starting out as superficial cells overlying the organizer and dorsal mesoderm at the beginning of gastrulation, and lining the dorsal posterior archenteron later-on (Blum et al., 2014). Interestingly, *pkd2* is involved at both stages as well in *Xenopus* (Vick et al., 2018); and data presented here). It remains to be seen to what extend this parallel holds, i.e. which other known molecules previously involved in either mesodermal process, participate in early LR axis formation as well. *brachyury* mutant mouse lines and homologous zebrafish *no tail* morphants support this reasoning, as do our unpublished results on the role of *brachyury* during LR development in *Xenopus* (Amack and Yost, 2004; King et al., 1998); Sabrina Kurz and MB, unpublished). In addition, mechanical strain has been recently shown to constitute a decisive physical force in LRO morphogenesis and function, namely cilia polarization and motility (Blum and Ott, 2018; Chien et al., 2018). Remarkably, strain is only effective when *foxj1* is present; therefore, it would be interesting to analyze whether or not FGF signaling is a prerequisite of strain-mediated LRO morphogenesis and function as well.

There is but one FGF-function in LR axis formation that remains unresolved in an evolutionary context: FGF-mediated vesicle release at the mouse LRO (posterior notochord or ventral node; (Tanaka et al., 2005). This is more than just a cursory inconsistency, as it impacts on our conceptional understanding and perception of cilia-mediated symmetry breaking in general. To this date, two models of symmetry breaking, though not necessarily mutually exclusive, coexist: (1) flow-sensing itself by immotile lateral sensory LRO cells, which through Polycystin-2 transmits a Ca^2+^ signal into the cells (2-cilia model; McGrath et al., 2003; Tabin and Vogan, 2003), and (2) the morphogen model, in which a secreted factor (a morphogen or vesicles; Nonaka et al., 1998; Okada et al., 1999), released from LRO cells, transfers through flow to the left side, where it initiates asymmetric signaling. In one variant of the morphogen model, the flow-transported factor is localized in vesicles (i.e. exosomes or exocytic vesicles), which are released in an FGF-dependent manner from the LRO. This variant, which suffers from a lack of follow-up studies since it was first proposed in 2005 has regained attention recently, following the proposal that sensory cilia do not qualify as calcium-responsive mechanosensors at the mouse LRO (Delling et al., 2016). This proposal contrasts with experimental and genetic evidence in the mouse, which demonstrated (a) that artificial flow was able to rescue lack of cilia motility as well as to revert laterality upon inversion of flow directionality (Nonaka et al., 2002); and (b) that Polycystin-2 is required on lateral LRO cells for flow sensing (McGrath et al., 2003; Yoshiba et al., 2012). The case, therefore, seems undecided at this point. In particular, it remains to be seen at what stage vesicles or a secreted morphogen arise at the LRO and how they transfer and fuse with cells or cilia on the left margin of the LRO. Our data, however, strongly suggest that the reported role of FGF in releasing vesicles at the mouse LRO is not conserved in the frog, as application of SU5402 after gastrulation did not impact on LR axis formation in our experiments. The LRO, from which such vesicles should emerge, only forms after gastrulation. Given the evolutionarily conserved functions of FGF signaling in mesoderm and LR development, our data seem to argue against a role of FGF in LRO vesicle release.

In conclusion, our work redefines the role of FGF signaling in vertebrate LR axis formation. In parallel to mesoderm induction, FGF specifies superficial organizer cells as LRO precursor during early gastrulation. This holds true despite apparent morphological differences between fish, amphibian and – perhaps – also mouse embryos. As mesodermal tissues undergo morphogenesis beginning at late gastrulation, FGF signaling, specifically the FGF/Ca^2+^ branch, is instrumental for the generation of flow sensor cells at the lateral margins of the LRO. LR axis formation, thus, seems to be more highly conserved as previously assumed. It remains to be seen whether this notion holds for flow sensing mechanisms as well.

## Materials and methods

### *Xenopus laevis* care and maintenance

Frogs were purchased from Nasco (901 Janesville Avenue P.O. Box 901 Fort Atkinson). Handling, care and experimental manipulations of animals was approved by the Regional Government Stuttgart, Germany (Vorhaben A379/12 ZO ‘Molekulare Embryologie’), according to German regulations and laws (§6, article 1, sentence 2, nr. 4 of the animal protection act). Animals were kept at the appropriate conditions (pH=7.7, 20°C) at a 12 h light cycle in the animal facility of the Institute of Zoology of the University of Hohenheim. Female frogs (4-15 years old) were stimulated with 25-75 units of human chorionic gonadotropin (hCG; Sigma), depending on weight and age, that was injected subcutaneously one week prior to oviposition. On the day prior to ovulation, female frogs were injected with 300-700 units of hCG (10-12h before). Eggs were collected into a petri dish by carefully squeezing of the females and *in vitro* fertilized. Sperm of male frogs was gained by dissection of the testes that was stored at 4°C in 1x MBSH (Modified Barth`s saline with HEPES) solution. Embryos were staged according to Nieuwkoop and Faber (1994).

### mRNA synthesis and microinjections

Prior to *in vitro* synthesis of mRNA using the Ambion sp6 message machine kit, the plasmid was linearized with NotI. Embryos were injected at the 4- to 8-cell stage, using a Harvard Apparatus. Drop size was calibrated to 8 nl / injection. Amounts of injected mRNA or MOs are indicated in the main text. For specific lineage targeting of constructs/MOs, tracers used for injection control included fluorescein (70,000 MW) and rhodamine B dextran (10,000 MW; both ThermoFisher). For more detailed lineage-specific injections to target central or lateral GRP and ventro-lateral tissues (LPM; cf. Blum et al., 2009a; Vick et al., 2018; Vick et al., 2009).

### Treatment with Fgfr1-inhibitor SU5402

SU5402 (Calbiochem) was diluted to 20mM in DMSO and stored in aliquots at −20°C. Incubations were conducted in 24-well plates and protected from light at room temperature. Concentrations used ranged from 30µM to 100µM, as indicated. Incubation start points and duration are indicated by NF stages in the main text. Incubations were terminated by removal of the inhibitor and several rounds of washes with buffer solution.

### In Situ Hybridization

For *in situ* mRNA detection, whole-mount in situ hybridization (ISH) was performed. Embryos were fixed in MEMFA for 2-3h and processed following standard protocols (Sive et al., 2000). RNA *in situ* probes were transcribed using SP6 or T7 polymerases. Whole-mount in situ hybridization was modified from Belo et al. (1997).

### Scanning electron microscopy and GRP analysis

Injected or treated specimens were fixed with 4% paraformaldehyde / 2.5% glutaraldehyde and processed for SEM analysis. SEM photographs were analyzed for ciliation, polarization and cell surface area by individual full GRP analysis using ImageJ and evaluated as described (Beyer et al., 2012; Sbalzarini and Koumoutsakos, 2005).

### Analysis of leftward fluid-flow

For analysis of leftward flow, dorsal posterior GRP-explants were dissected from control or SU5402-treated embryos at stage 17. GRP-explants were placed in a petri-dish containing fluorescent microbeads (diameter 0.5 µm; diluted 1:2500 in 1xMBSH) and incubated for a few seconds. Explants were transferred to a microscope slide which was prepared with vacuum grease to create a small chamber that contained fluorescent microbeads solution; a cover slip was carefully pressed on to seal the chamber. Time lapse movies of leftward flow were recorded using an AxioCam HSm video camera (Zeiss) at 2 frames per second using an Axioplan2 imaging microscope (Zeiss). For flow analysis, ImageJ and statistical-R were used. Using the Particle-Tracker plug-in from ImageJ, leftward flow was analyzed, particle movement was measured, and data were processed as described previously (Beyer et al., 2012; Sbalzarini and Koumoutsakos, 2005; Vick et al., 2009).

### GRP width analyses

For relative quantification of GRP width, either SEM photographs of broken GRPs or vibratome sections of *myod1*-stained embryos were used. For evaluation of *myod1* expression, sections containing the highest number of *myod1*-positive cells were selected. Width was measured in pixels using ImageJ (Sbalzarini and Koumoutsakos, 2005). For each analysis, the mean value of DMSO-treated controls was set to 1.0 and the relative sizes of manipulated embryos was calculated to generate box plots.

### Statistical analysis

Statistical calculations of marker gene expression patterns and cilia distribution were performed using Pearson’s chi-square test (Bonferroni corrected). For statistical calculation of ciliation, cilia length, cell size, flow velocity and directionality Wilcoxon-Match-Pair test was used (statistical R).

*=p<0.05, **=p<0.01, ***=p<0.001 were used for all statistical analyses.

## Acknowledgements

The authors like to thank especially Thomas Thumberger for expert help with flow analysis, Tina Beyer for technical advice, and E. Amaya, C. Kintner, R. Rupp, H. Steinbeisser, and A. Vonica for reagents.

